# UAV Based Imaging Platform for Monitoring Maize Growth Throughout Development

**DOI:** 10.1101/794057

**Authors:** Sara B. Tirado, Candice N. Hirsch, Nathan M. Springer

## Abstract

Plant height (PH) data collected at high temporal resolutions can give insight into important growth parameters useful for identifying elite material in plant breeding programs and developing management guidelines in production settings. However, in order to increase the temporal resolution of PH data collection, more robust, rapid and low-cost methods are needed to evaluate field plots than those currently available. Due to their low cost and high functionality, unmanned aerial vehicles (UAVs) can be an efficient means for collecting height at various stages throughout development. We have developed a procedure for utilizing structure from motion algorithms to collect PH from RGB drone imagery and have used this platform to characterize a yield trial consisting of 24 maize hybrids planted in replicate under two dates and three planting densities in St Paul, MN in the summer of 2018. The field was imaged weekly after planting using a DJI Phantom 4 Advanced drone to extract PH and hand measurements were collected following aerial imaging of the field. In this work, we test the error in UAV PH measurements and compare it to the error obtained within manually acquired PH measurements. We also propose a method for improving the correspondence of manual and UAV measured height and evaluate the utility of using UAV obtained PH data for assessing growth of maize genotypes and for estimating end-season height.

## INTRODUCTION

Plant height (PH) serves as a major growth indicator and can be used for assessing crop productivity and making crop management decisions. PH has been shown to be linked to nitrogen (N) nutrition during vegetative development in maize (Yin et al., 2011-1), making it useful for assessing spatial variability in crop response to N. PH can therefore help guide precision agriculture practices such as variable-rate N applications within the field. PH has also been linked to plant biomass in maize and barley crops (Li et al., 2016, Bendig et al., 2014). Studies have found PH at early to mid-developmental stages in maize are correlated to grain yield (Yin et al., 2011-2; Katsvairo, Cox and Van Es, 2003), and can be useful for forecasting crop yields (Yin et al., 2011-1). Thus, tracking plant height at these earlier stages and throughout development can help to identify superior cultivars in plant breeding programs and for developing management practices that account for spatial heterogeneity in production fields. To maximize the utility of PH data, height at various stages throughout development needs to be gathered, and with current methods for measuring height this can be challenging. Similar to many phenotypic traits, current practices of gathering height in field settings involve physically measuring PH with a large ruler, which is time consuming and difficult to implement on a large scale. Ruler measurements are also subject to user bias and error, decreasing the accuracy and utility of these measurements.

Remote sensing technology has been proposed as an efficient means for collecting rapid and objective measurements for PH (Madec et al., 2017, Yang et al., 2017). Unmanned aerial vehicles (UAVs) have proven to be a promising platform for collecting remote sensing data as they are inexpensive yet capable of achieving very high spatial and temporal resolutions. Many modern UAVs also have the functionality of flying automatically along a path specified by the user through mission planning software, increasing reproducibility through time. Commercial UAV platforms come equipped with RGB cameras that can be utilized to capture images along the field to create 3D field reconstructions using structure from motion (SfM) and multiview stereo (MVS) algorithms which rely on estimating the 3D structure of a scene utilizing a set of 2D images (James and Robson, 2012). The speed and ease with which UAVs can be flown for data acquisition provide advantages in scale and temporal resolution over other remote sensing methods utilized for creating 3D models, such as light detection and ranging (LiDAR), which involve high costs in data acquisition and processing (Díaz-Varela et al., 2015). However, there is substantial improvement that is needed in developing tools and approaches for simple implementation of UAV technology to generating phenotypic data in field settings.

There are examples of success in using RGB sensors to extract a number of traits that, if taken through time, can give us insight into plant growth and the relative effects of genetic and environmental variables on these traits. Canopy coverage estimates have been gathered for soybeans using digital cameras and these have been found to highly correlate to canopy light interception measurements (Purcell, 2000) and grain yield (Xavier et al., 2017). Previous studies have estimated PH from UAV imagery in sorghum (Chang et al, 2017; Shi et al., 2016; Watanebe et al., 2017), wheat (Madec et al., 2017; Michalski et al., 2018; Holman et al., 2016), cotton (Feng et al., 2018), barley (Bendig et al., 2014) and maize (Anderson II et al., 2019; Anthony et al., 2014; Geipel, Link and Claupein (2014); Grenzdörffer, 2014; Su et al., 2019; Varela et al., 2017). Although these studies have reported high correlation in PH measurements across various dates to manual measurements, they lack estimates of how daily correlations of imagery-derived PH (PH_UAV_) and ruler-derived PH (PH_R_) vary throughout different growth stages and how this compares to the inherent error in PH_R_ measurements.

Taking into account past efforts in height estimation from UAV platforms, the present work aims to implement SfM algorithms for the estimation of PH_UAV_ for a maize field throughout development and compare these measurements to daily PH_R_ measurements collected using the traditional manual ruler-based method. We tested the error in PH_R_ measurements and assessed the correlation of PH_UAV_ to PH_R_ measurements. We also evaluated whether models developed using PH_UAV_ measurements together with PH_R_ measurements of only a small subset of plots could improve the correspondence of mean plot PH_UAV_ measurements to mean plot PH_R_ measurements.

## MATERIALS AND METHODS

### Experimental Design

Two replicates of 12 maize hybrids were planted in 4-row plots at two dates (05/14/2018 and 05/21/2018) and three densities (60k, 90k and 120k plants per hectare) following a Randomized Complete Block Design blocked by planting date and replicate within planting date in St Paul, MN in the summer of 2018 (Figure S1). Each row was 15ft long center-to-center with 12 ft of plots and 3ft of alleys and 30in spacing between rows. There were a total of 144 four-row plots planted for this experiment. The 12 hybrid genotypes utilized were generated by crossing ex-PVP lines selected due to their past use in production settings and the availability of seed. This experiment was grown on two acres. Nine 1m x 1m ground targets were distributed around the border and internal alleys of the field for use as ground control points (GCPs) based on previously developed optimization algorithms (Gomez-Candon, De Castro and Granados, 2014).

### Drone Image Data Collection

The field was imaged approximately weekly from planting until plants reached terminal height using a DJI Phantom 4 Advanced drone flown 30m above ground to achieve a ground sampling distance (GSD), measured as the ground distance represented between the centers of two neighboring pixels, of approximately 0.82cm. SfM algorithms for 3D reconstruction of field points from 2D images rely on finding correspondences between images. Wind can cause issues by causing the object to move between two consecutive recordings. This can be reduced by using a high measuring repetition rate (Paulus, 2019). Images were collected in a grid pattern using 85% frontlap and 85% sidelap to maximize reconstruction efficiency (Figure 1B). A total of 615 images were gathered per mission and a total of twelve missions were conducted throughout the season.

**Figure 1.**
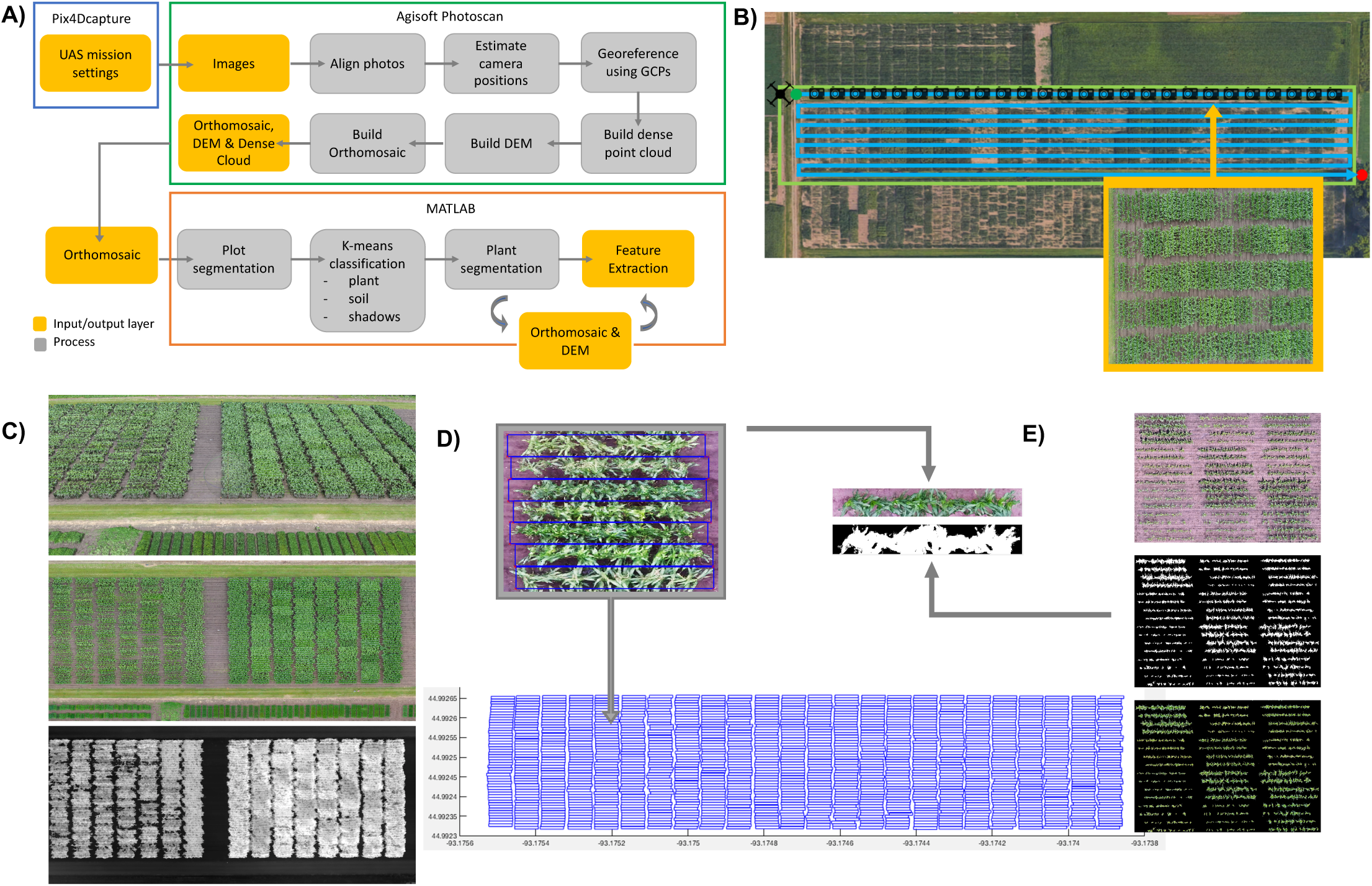
Procedure for feature extraction from UAV images. **A)** Image generation and processing pipeline. **B)** UAS flight mission structure for gathering images. **C)** Dense cloud (top), orthomosaic(middle) and DEM (bottom) output from Agisoft Photoscan Professional Edition. **D)** Plot boundary extraction from grid shapefile for plot segmentation. **E)** Plant material segmentation using k-means algorithm. The original image (top), plant segmentation mask (middle) and applied mask on original image (bottom) are shown.

### Manual Measurement Data Collection

Manual measurements for plant height (PH_R_) were collected on the same day as eight of the drone flights from two plants per row from each of the middle two rows of each four-row plot using a ruler. PH_R_ was measured as the distance between the ground and the topmost freestanding vegetative part of the plant until plants reached reproductive growth and then to the top point of the tassel. For 18 selected plots, hand measurements of all plants in one of the middle two rows were collected to get estimates of variance in height within a row. The 18 plots comprised of one replicate of all three densities in the early planting date for 6 randomly selected genotypes (DK3IIH6 x LH198, LH198 x PHN46, LH82 x PHK76, PHB47 x LH198, PHB47 x PHP02 and PHK56 x PHK76).

### Data Processing Workflow

Agisoft software (Agisoft Photoscan Professional Edition v1.4.4, 2018) was used to process the collected imagery and generate crop surface models (CSM) and RGB orthomosaics for each date following the developed workflow (Figure 1A). Processing steps implemented in Photoscan include feature matching, solving for camera intrinsic and extrinsic orientation parameters, building a dense point cloud, building a field orthomosaic, and building a digital elevation model as specified by Photoscan manual recommendations (Agisoft LLC, 2018; Figure 1C). Parameters set for image alignment included: high accuracy for obtaining camera position estimates, reference pair selection so that overlapping pairs of photos are selected based on measured camera locations, a key point limit of 40,000 points, and a tie point limit of 4,000 points. Ground control points were then used to optimize camera alignment. For generating the dense point cloud, the quality value was set to High with Moderate depth filtering to optimize processing time and model quality.

QGIS software (QGIS v2.18.9, 2017) was utilized for plot boundary extraction by overlaying a grid based on plot size and spacing and exporting plot boundary coordinates for the middle two rows of each plot (Figure 1D). Custom MATLAB algorithms were developed to process the CSMs and orthomosaics to extract height estimates for individual plots. Plant height was extracted by segmenting individual plots from the CSMs, overlaying a grid with 20 bins along the plot and extracting the 3^rd^ percentile and 97^th^ percentile height values for each cell. The ground height for the plot was defined as the minimum 3^rd^ percentile of all cells in the plot as this value represents a ground height value in the alley. To extract the PH_UAV_ value for each cell, the ground height value (minimum 3^rd^ percentile across cells) was subtracted from the 97^th^percentile height value in each cell. To calculate the mean PH_UAV_ per plot, a trimmed mean function was implemented on the PH_UAV_ values of the middle 12 bins of the plot trimming the two extremes and getting the average of the remaining 10 values (Figure 2). Only the middle 12 bins for the middle two rows in each plot were assessed when calculating an average PH value to prevent the average height for a given plot to be biased by the height of plants near alleys or different density treatments. A piecewise polynomial spline of order 20 was fit using the MATLAB splinefit function to better represent the continuous plant growth that occurs throughout the growing season (Lundgren, 2019). To assess how growth changes through time, we then extracted the first derivative, or slope, of the curve to get change in height through time.

**Figure 2.**
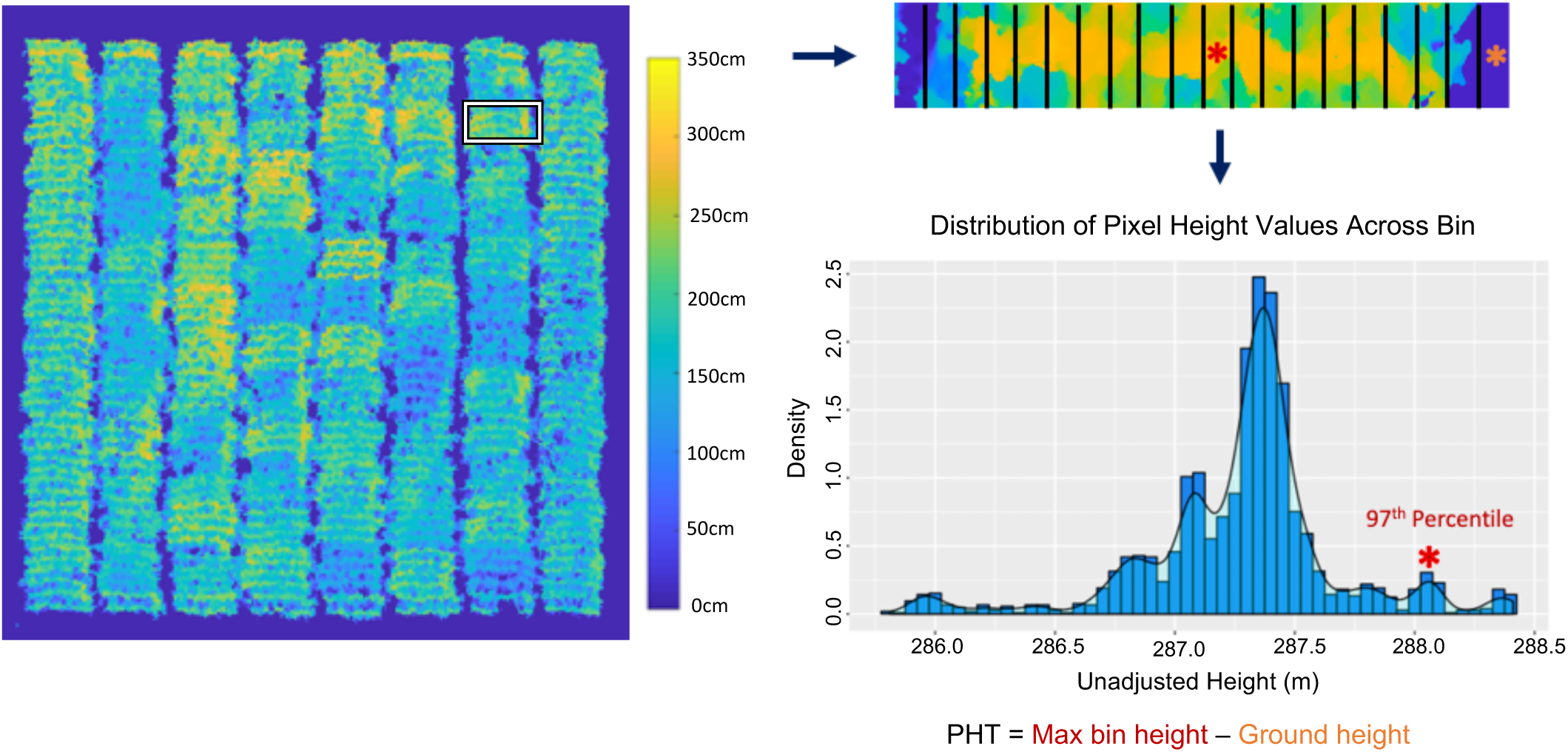
Method for extracting mean PH_UAV_ values for a given plot. The plot is segmented and broken down into 20 bins. The 3rd and 97th percentiles for each bin are extracted and used to calculate the average plot height by subtracting the minimum 3rd percentile of all bins from the 97th percentile of each bin.

### Statistical Analyses

A total of 1,152 mean plot PH_UAV_ values were extracted across eight timepoints. Outliers were identified for each timepoint as those values that were less than or larger than the mean PH_UAV_ value for that date plus or minus the standard deviation multiplied by a factor of 2½. A total of 17 outlier points across all dates were identified and removed from the dataset for subsequent analysis. It should be noted that while there was a strong wind event that caused substantial lodging in the experiment (see below), these outlier points were random and not enriched for data points collected at or shortly after the lodging event. To measure the strength of the linear relationship between different PH measurements, Pearson’s correlation was utilized.

Variation in plant height values was partitioned into fixed effects for genotype, planting date, density, genotype-by-density interaction, and genotype-by-planting date interaction and random effect of replicate with a linear mixed model using the lmer function of the lme4 R package (Bates et al., 2015). To test the significance of the effects of the different variables of the mixed linear model on plant height, we used the Anova function of the R car package to run a Type II Wald chi-square test (Fox and Weisberg, 2019).

### Model Selection

A model was built for each time point using PH_UAV_ values along with other explanatory variables to better predict PH_R_ values for a given date. An optimal model was selected at each time point based on an original model containing the variables of PH_UAV_ measurements on that date, genotype, planting date, and categorical planting density using a backward stepwise model with the step function of the stats R package which conducted model selection by AIC (R Core Team, 2012). These predictor variables were chosen based on their biological foundation as well as their level of significance for predicting PH_R_ values. The selected model in every timepoint contained the variables PH_UAV_ measurements, genotype, and planting date. However, the model coefficients for each variable were specific to each timepoint.

The scripts and processes used to perform the image analyses and trait extract are available at https://github.com/SBTirado/UAV_PH.git.

## RESULTS AND DISCUSSION

Our goal was to develop a robust low-cost platform for efficient estimation of plant height for maize plots. High-throughput collection of plant height measurements throughout the growing season can provide a better understanding of how different environments influence maize growth and can help to dissect the underlying cause of end of season genotype x environment (GxE) interactions that are commonly observed. In order to develop a robust plant height collection pipeline and to begin to consider the factors that influence plant growth rates, we planted a set of 12 hybrids in a design that incorporated 3 planting densities of 60K, 90k or 120k plants per hectare and two planting dates (Figure S1). There were two replicate plots of each genotype at each planting density and this full set of materials (2 replicates of 3 planting densities) was planted at two different dates such that plants would be exposed to similar environments but at different growth stages. To decrease the impact of neighboring plots, each plot was grown as a four-row plot and only plants in the middle two rows were measured. Manual measurements were taken using a ruler (PH_R_) for two middle plants from the two rows in the middle of the 4-row plot for each plot on the dates that aerial images were obtained. Analysis of variance showed significant variation in final plant height within this experimental design due to genotype, planting date, and density (Table 1).

**Table 1.** Analysis of variance of hand measured terminal plant height for 12 hybrids planted in a randomized block design with density and planting date treatments.

|  |  |  |  |  |
| --- | --- | --- | --- | --- |
|  | **Degrees of Freedom** | **Sum of Squares** | **Mean Sum of Squares** | **Percent of total variation** |
| Planting Date | 1 | 31241 | 31241 | 34.66*** |
| Genotype | 11 | 40518 | 3683 | 44.95*** |
| Density | 2 | 764 | 382 | 0.85* |
| Planting Date:Genotype | 11 | 7194 | 654 | 7.98*** |
| Density:Genotype | 22 | 2042 | 93 | N.S. |
| Residuals | 96 | 8374 | 87 | N.S. |
| * significant at $p=0.05$; ** significant at $p=0.01$; *** significant at $p=0.001$; N.S., not significant. | | | | |

Aerial images of the field were collected on eight different dates throughout the growing season until plants reached terminal height (approximately weekly, weather permitting). At each date there were ∼615 images collected of the field with 85% frontlap and sidelap between images. The images were used to create high resolution 3D models, field orthomosaics, and DEMs of high resolution using existing SfM algorithms (Figure 1). There are multiple methods that have been utilized for extracting plant height from UAV images. A common approach used is the difference based method (DBM) which involves subtracting a digital terrain model (DTM) of the bare ground from the DEM (Varela et al., 2017). The DTM can be generated by flying the bare ground but requires an additional day of data collection and processing and can introduce error due to minor fluctuations in flight altitude within and across dates that are common with most commercial drone platforms. Other approaches attempt to create the DTM by identifying and extracting ground points from the pointcloud and interpolating the height of intermediate points to create a surface model of the ground and then taking the difference between the DTM and DEM, but these approaches require processing large 3D point clouds and finding optimal parameters for the interpolation algorithm which can be computationally intensive (Su et al., 2019). Different algorithms for generating a DTM including DBM and three point cloud interpolation methods were evaluated by Anderson II et al. (2019) to see which provides higher accuracy when estimating plot heights. They found that all methods had similar, consistent performance in flat, uniform fields like those used in breeding trials.

To account for the drawbacks and challenges of the other proposed methods and because breeding field trials are typically in flat and uniform fields, rather than generating a DTM, we developed a pipeline that involves identifying ground height values individually for each plot when calculating mean plot height. For each plot the height was estimated from the UAV imagery (PH_UAV_) by calculating the difference between the ground height and surface height of plant material across each plot and calculating a mean plot height value (Figure 2, see methods). Although not analyzed in this study, this method also allows for the calculation of variance in PH_UAV_ throughout a plot.

### Error within ruler height measurements

A critical question for any application using PH_UAV_ is the quality of the height estimates that are obtained through this high throughput measurement. However, there are multiple complicating factors in truly assessing the quality of PH_UAV_ estimates. An important factor is that the manual measurements of plant height (PH_R_) that are used as the ‘ground truth’ for determining the accuracy of PH_UAV_ estimates can include some level of error. In order to begin to understand the quality of the PH_R_ estimates, we replicated hand measurements of the same plants for a subset of plots at three of the timepoints (6/13/18, 7/17/18, and 8/9/18). To do this, we measured the same two plants per row twice for 12 randomly selected plots by having personnel return to the same plots when taking manual measurements and measuring the same plants which were tagged. An assessment of the correlation and RMSE in replicated hand measurements of the individual plants and plot mean PH_R_ revealed variation in the correlations and RMSEs (Table S1). In the earliest timepoint which corresponds to crop vegetative stages of V6 and V4 for the first and second planting respectively, the correlation between ruler measurement replicates showed the lowest correlation (adj. r-square value of 0.45 for individual plant measurements and 0.64 for plot mean measurements calculated using the two plants from the middle two rows). This correlation improved with later developmental stages. On the other hand, the RMSE between ruler measurement replicates was low in the first timepoint but remained fairly constant afterwards. The correlation at terminal plant height was high (adj. r-square value of 0.9 for individual plant measurements and 0.64 for plot mean measurements) and had a low amount of error (RMSE of 7.18cm for individual plant measurements and 3.00cm for plot mean measurements; Figure 3B).

**Figure 3.**
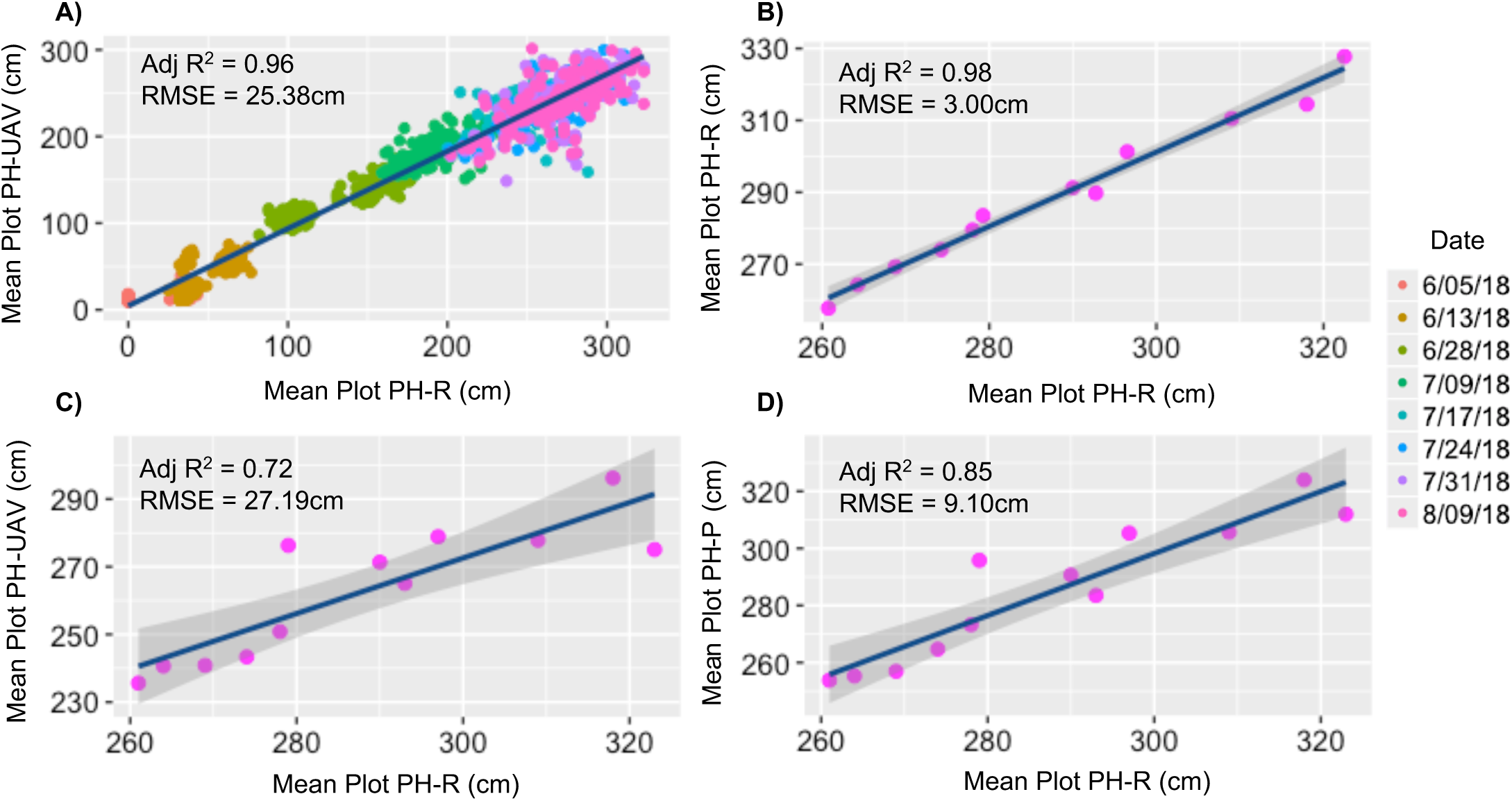
Pearson correlation plots of various PH measurements. **A)** Correlation of mean plot PH_UAV_ and mean plot PH_R_ across all dates. **B)** Correlation of mean plot PH_UAV_ and mean plot PH_R_ for a subset of plots that was hand measured twice in 8/09/2018. **C)** Correlation of mean plot PH_R_ for hand-measurement replicates of a subset of plots were same plants were hand measured twice in 8/09/2018. **D)** Correlation of model-estimated PH_P_ and mean plot PH_R_ for a subset of plots that was hand measured twice in 8/09/2018. PH_P_ was estimated using a linear regression model that included PH_UAV_ and variables that contributed to observed variation in height values.

Another potential difference between PH_R_ and PH_UAV_ estimates of plot height is that manual measurements are often collected for a subset of plants in a row while the UAV measurements are often generated as a per plot mean height that samples a larger number of plants. To investigate the accuracy in plot mean PH_R_ obtained by measuring only two random plants per row, we randomly sampled two individual plant PH_R_ measurements at each timepoint twice for the 17 plots in which all plants for one of the middle two rows were measured and calculated new replicates for plot mean PH_R_ measurements with the intention of assessing the accuracy and error in traditional methods of plot mean calculation. We replicated this procedure 100 times for each plot and timepoint to get an estimate of the average amount of error in PH_R_ mean values calculated by measuring only two plants per row. Plot mean PH_R_ measurements obtained from different sets of plants within a row showed a much lower correlation and larger error across most timepoints compared to the correlations between PH_R_ replicated measurements of the same two plants (Figure 4D; Table S1). These correlations are comparable to those obtained when correlating plot mean PH_UAV_ measurements to plot mean PH_R_ measurements (Figure 4A). This indicates that PH_R_ measurements can have substantial variation that is impacted by the number of plants measured and the selection of plants to be measured.

**Figure 4.**
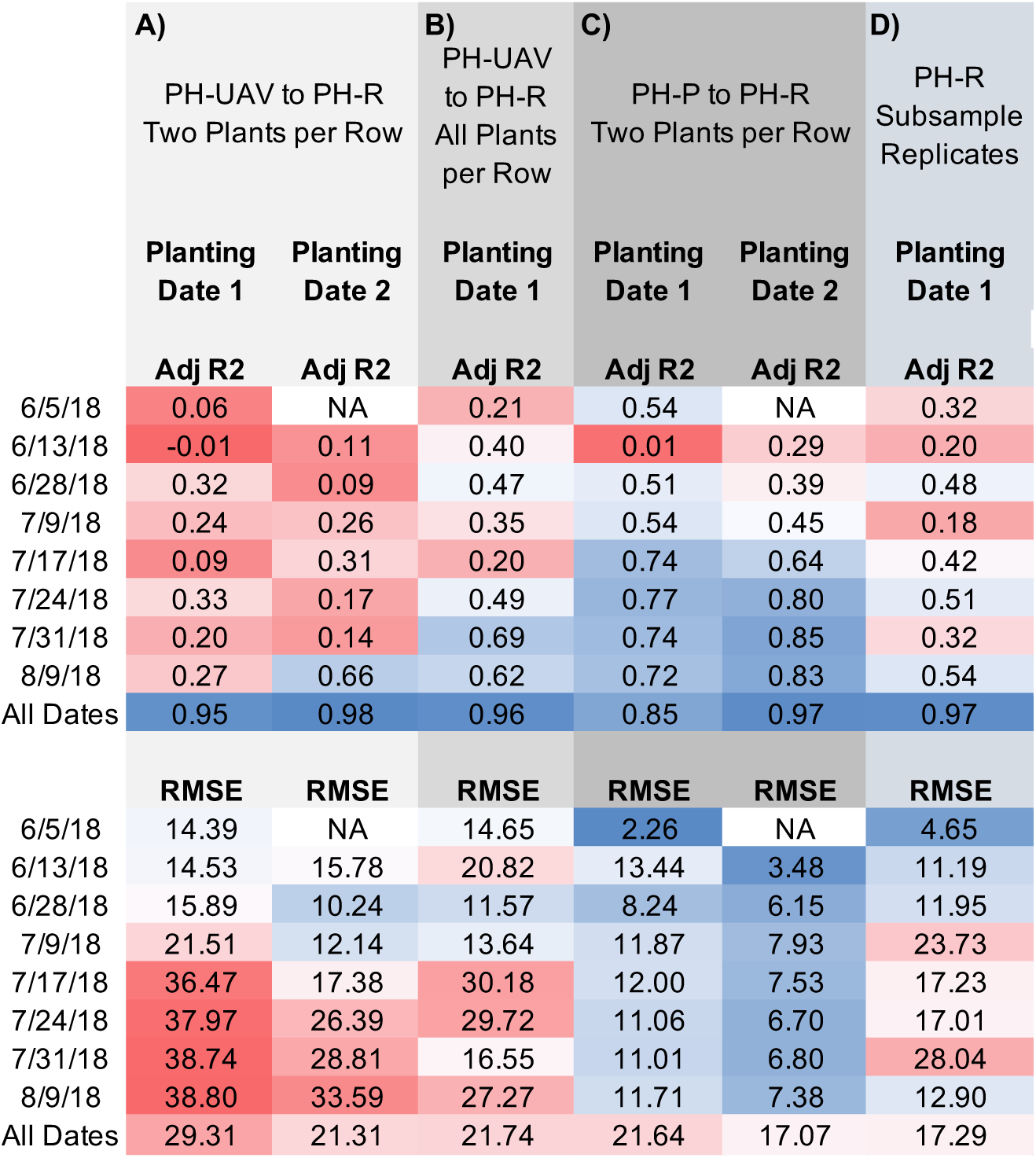
Adjusted r-square values and root mean square error for the linear correlation of various PH measurements. **A)** UAV-derived (PH_UAV_) plot mean height values compared to the respective hand-measured plot mean height (PH_R_) for plots where two plants per row were hand measured. **B)** UAV-derived (PH_UAV_) plot mean height values compared to the respective hand-measured plot mean height (PH_R_) for plots where all plants for one of the middle two rows were hand measured. **C)** Model-derived (PH_P_) plot mean height values using UAV height measurements compared the respective hand-measured plot mean height values (PH_R_) **D)** Means derived from two iterations of random sampling of hand-measured height values for two plants across plots were all plants for one of the middle two rows were hand measured.

### Correlation of ruler height to algorithm height measurements

There are limitations in accuracy of ruler-estimated heights for specific plants as well as for estimating the height of a plot. However, these measurements can still be used to assess the quality of PH_UAV_ measurements, if we accept that the quality of the correlations between PH_UAV_ and PH_R_ likely cannot be improved beyond the inherent errors in PH_R_ measurements. Comparison of PH_R_ and PH_UAV_ showed the overall correlation across all data collection dates was strong (adj. r-square value of 0.96 for our dataset; Figure 3a), as has been previously shown in other studies (Varela et al., 2017). However, the correlations within a single date were much lower (Figure 4A). This is likely due to the fact that the level of variation for height on a single date tends to be relatively low. The root mean square error (RMSE) for mean plot PH_UAV_ compared to the mean plot PH_R_ across all dates and both planting dates was 25.38cm, but varied from 14.39cm to 38.80cm across dates for the first planting date and from 10.24cm to 33.59cm across dates for the second planting date, with a general trend of an increasing RMSE with growing developmental stage (Table S1). The lower correlations and higher RMSE for the first planting date was likely due to a storm that occurred on July 1st that reached a peak wind speed of 50 mph (National Weather Service, 2018). This storm caused large degrees of lodging, especially within plots in the early planting date treatment causing larger variability in height among plants within plots.

If we compare the error in PH_UAV_ measurements to that in PH_R_ replicated measurements for the subset of plots where two individual plants were hand-measured twice, we see that the correlations of PH_R_ and PH_UAV_ measurements for the same plots increased throughout development but were lower than the correlation between PH_R_ replicated measurements (Table S1). By terminal height the correlation between PH_R_ and PH_UAV_ measurements was high (adj. r-square value of 0.72) but had a substantial amount of error (Figure 3C). Unlike the high correlation and low RMSE obtained from plot mean PH_R_ measurement replicates calculated by measuring the same two plants per row, plot mean PH_R_ measurements obtained by sampling different sets of plants within a row showed a much lower correlation and larger error across most timepoints (Figure 4D; Table S1). These correlations between plot mean PH_R_ calculated by measuring varying plants in a row, however, are comparable to those obtained when correlating plot mean PH_UAV_ measurements to plot mean PH_R_ measurements (Figure 4A). The direct comparison of the PH_R_ and PH_UAV_ measurements obtained therefore yielded an error comparable to the inherent error in estimating PH_R_ across most timepoints.

Given that mean plot PH_UAV_ estimates take into account variation in height throughout a plot, we can better assess their correlations to mean plot PH_R_ measurements by looking at the subset of plots where all plants in a row were measured. Doing so, we see that the correlations and RMSE estimates greatly improve relative to those calculated by comparing mean plot PH_UAV_ estimates to mean plot PH_R_ measurements estimated by measuring only two plants per row (Figure 4B). This indicates that mean plot PH_UAV_ measurements better resemble the true plot mean compared to the mean calculated from measurements of just a subset of plants per row.

### UAV-estimated heights can detect biological variation

Our goal was to develop a low cost, high-throughput platform for monitoring PH through time that would allow us to look at how genotypes develop and respond to different environmental conditions. A storm with wind speeds reaching a peak of 50 mph occurred on July 1st of the 2018 growing season in St. Paul, providing a prime opportunity to evaluate the utility of this method to track plant responsiveness to environmental conditions (National Weather Service, 2018). An example of this responsiveness can be seen in the height profiles for LH82 x DK3IIH6 plots, where plants at high density in the earlier planting date showed a reduction in height 45 days after sowing (Figure 5A). Normally, a reduction in height would be unexpected and may reveal issues with the height estimation platform, but this height reduction was due to the lodging that occurred as a result of the storm. To look at the lodging event more broadly, we calculated the percent change in height relative to the last images collected three days prior to the storm (Figure 6). This revealed substantial differences in percent and degree of lodging that were influenced by several factors. As with the example genotype (Figure 5A), there was very little lodging in plots from the second planting date, likely because these plants were smaller at the time of the storm and were minimally affected (Figure 6). There were also differences in the amount and severity of lodging for the three planting densities (Figure 6). UAV imagery provides an opportunity to collect data rapidly in response to events such as this storm event and track plants at regular intervals that is much more difficult to do with hand measurements. Indeed, we were not able to track the event in the PH_R_ measurements (Figure 5B) as these measurements were not taken until several days after the wind storm and most plots recovered from the lodging prior to this measurement.

**Figure 5.**
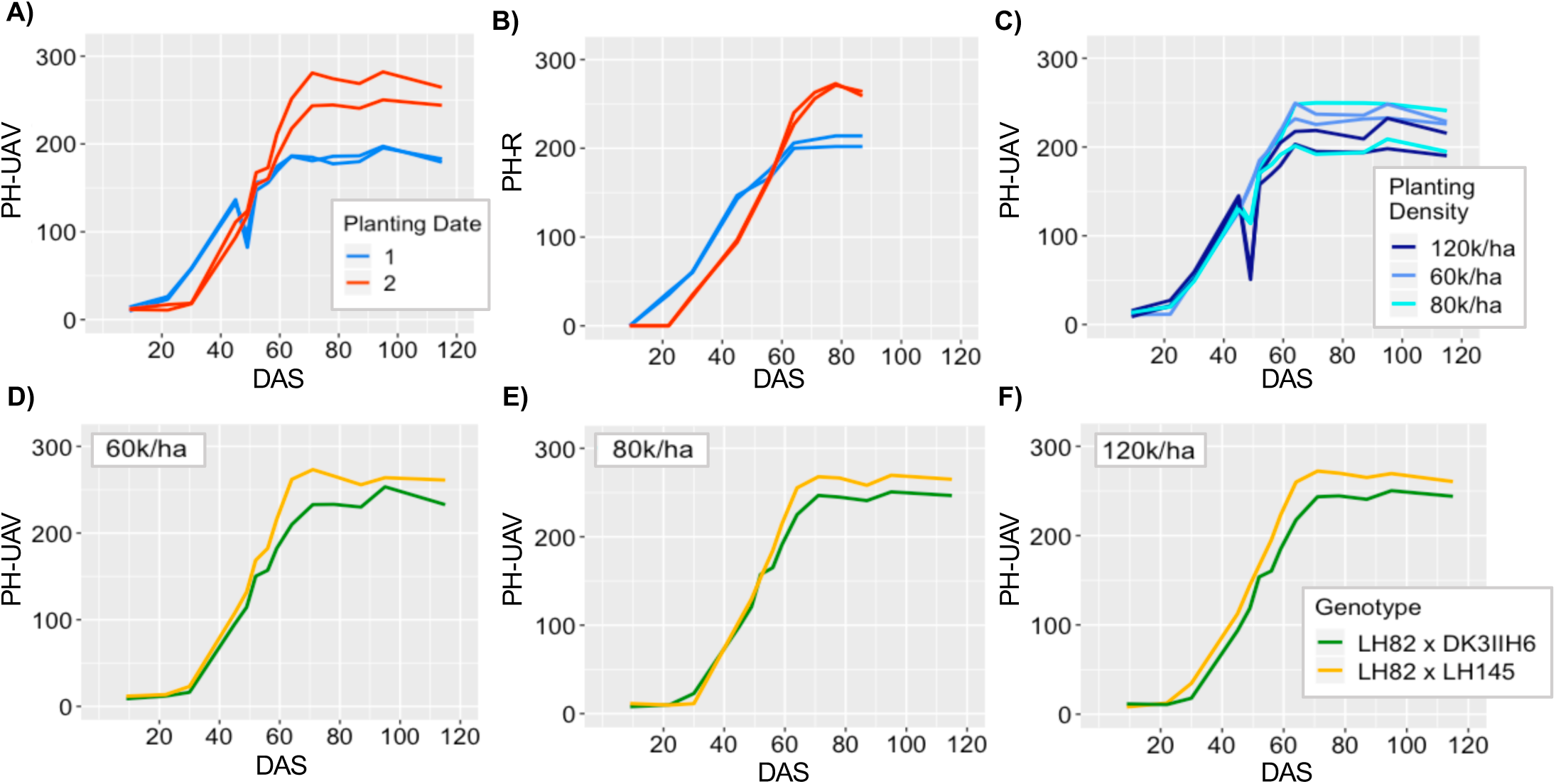
Height through time for various genotypes and treatments. **A-B)** Height through time as measured by the UAV (A) and hand measurements (B) for two replicate plots of a single genotype (LH82 x DK3IIH6) planted under high density across two planting dates. **C)** Height through time as measured by the UAV for two replicate plots of a single genotype (LH82 x PHK76) in the early planting date treatment and the three planting densities. **D-F)** Height through time as measured by the UAV for plots of two different genotypes (LH82 x DK3IIH6 and LH82 x LH145) in the late planting date treatment and the low (D), medium (E) and high (F) planting densities.

**Figure 6.**
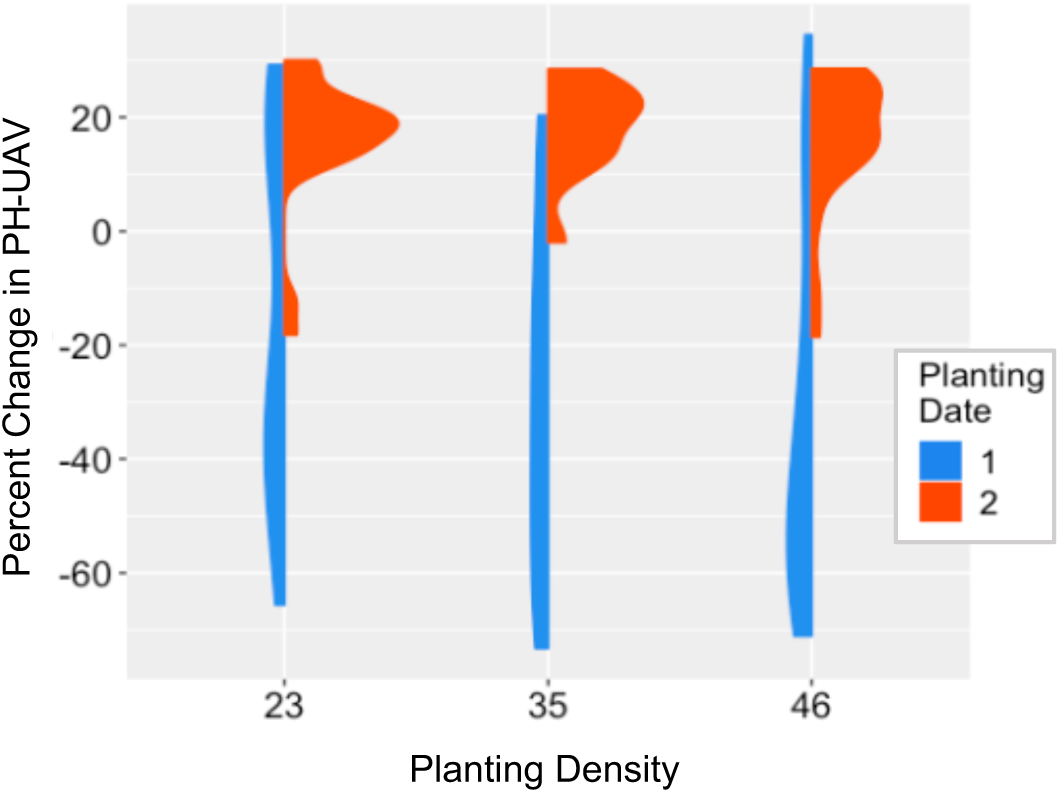
Percent change in UAV mean plot height from 6/28/18 to 7/2/18 for all plots across both planting dates. Change in height was due to the storm that occurred on the night of 7/1/18.

The PH_UAV_ estimates were also able to document differences in growth rates throughout the growing season between genotypes. For example, LH82 x LH145 planted at high density in the late planting had a faster rate of growth compared to LH82 x DK3IIH6 planted in the same conditions as soon as 20 days after sowing (Figure 5F). LH82 x LH145 was able to maintain an advantage in terms of height throughout development and achieve a higher realized height at the end of the season. The ability to track these differences in growth patterns among genotypes that may or may not impact the end of season heights that are typically collected provides additional information to breeders in the process of selecting superior parents and hybrid combinations.

We expected to note differences in height within genotypes and planting dates based on planting density. Plants grown at higher density tend to grow faster and taller when water and nutrient resources are not limited as they are competing with more plants for sunlight (Tetio-Kagho & Gardner, 1988). However, our data suggests that the realized height remains pretty stable across densities and that in many cases, the differences in height due to genotypic variation were reproducible across planting dates and densities (Figure 5D-F). Although density had only a small impact on end-season height, it did contribute to differences observed for lodging across genotypes in the early planting date. Plots planted at higher densities experienced a more severe degree of lodging when exposed to strong winds compared to plots planted at a lower density (Figure 5C). Overall, visualization of the plant height profiles for each genotype x condition suggested a fairly robust ability to monitor biological variation for height and responsiveness to environmental conditions under different growth conditions.

### Creating a model to improve height estimation accuracy

Given that ruler-derived height is the standard way of collecting PH measurements, we wanted to find a procedure that could improve the correlations between PH_UAV_ and PH_R_ measurements. We hypothesized that using information about height collected for a subset of individuals together with factors that influence plant morphology and height could be used to improve the accuracy of estimated height from UAV images and therefore the correlation with PH_R_ measurements. In order to test this hypothesis, we assessed whether creating a model based on hand measurements of a subset of plots could improve the correspondence of PH_UAV_ estimates to PH_R_. An optimal model based on a set of predictor variables was selected at each time point using a backward stepwise model followed by model selection by AIC (see methods). The selected model for all timepoints incorporated mean PH_UAV_ values for each plot as a predictor variable together with factors that were shown to contribute to the variation observed for hand height measurements including the plot genotype and planting date. A linear regression model was derived from 1/3 of the dataset following model selection to generate predictions (PH_P_) on the remaining plots.

Correlation between the PH_P_ estimates and measured height values were substantially higher using the derived model compared to the direct use of mean plot PH_UAV_, and the correlations improved significantly after the V8 crop developmental stage (Figure 4C). Moreover, the RMSE for the estimated height values compared to the hand measurements improved and ranged from 2.26cm to 13.44cm in the early planting and from 3.48cm to 7.93cm in the late planting. These error estimates are about half the size of the error when comparing straight PH_UAV_ to PH_R_ measurements. The same trend can be seen for plots where hand-measurement replicates were obtained (Table S1). At terminal height, the correlation of model-derived PH estimates to mean plot PH_R_ values for these plots was higher than when comparing PH_UAV_ estimates, and the RMSE was substantially lower (Figure 3D). Sampling a portion of the field plots to get estimates of height that closely correlate to what we would get with traditional methods for height estimation saves time and labor costs when a large number of plots are involved. Given that PH_R_ measurements are the standard way of collecting plant height data, having PH_UAV_ measurements that correlate is necessary for being able to compare height for genotypes grown across different locations and multiple years to historical data and contemporary data collected in other groups not using aerial imagery.

### Predicting end-season height from height collected at earlier time points

Being able to estimate height for any given date is useful for tracking growth; however, being able to predict end-season height using height from earlier time points can allow for the assessment of end-season performance at a time when you can still implement management practices to enhance it. This also can allow breeders to make selections of top-performing plants that meet the desired phenotypic characteristics before making crosses thereby helping maximize their selection efficiency. We wanted to test whether height at early developmental stages could be predictive of terminal height. A previous study by Li et al. (2018) showed low correlations between PH collected for hybrid varieties at early developmental stages to their respective end of season height likely due to differences in the physiological basis for PH at both stages. PH early in development is driven primarily by genetic factors; however, environmental factors begin to impact terminal height achieved as the plant develops. Including temporal data for height, or rather the rate of change in height through time, could be more informative when predicting terminal height as it captures how different genotypes interact with the environment and how this interaction impacts the trait at hand. Therefore, rather than just using height at specific timepoints (Figure 7A) to predict end season height, we wanted to utilize rate of growth at intervals in time. To do this, a B-spline, or piecewise polynomial function, of 20 degrees was fit to create a smooth curve representative of true growth. The first derivative, or slope, of the B-spline curve was then extracted to get change in height through time (Figure 7B).

**Figure 7.**
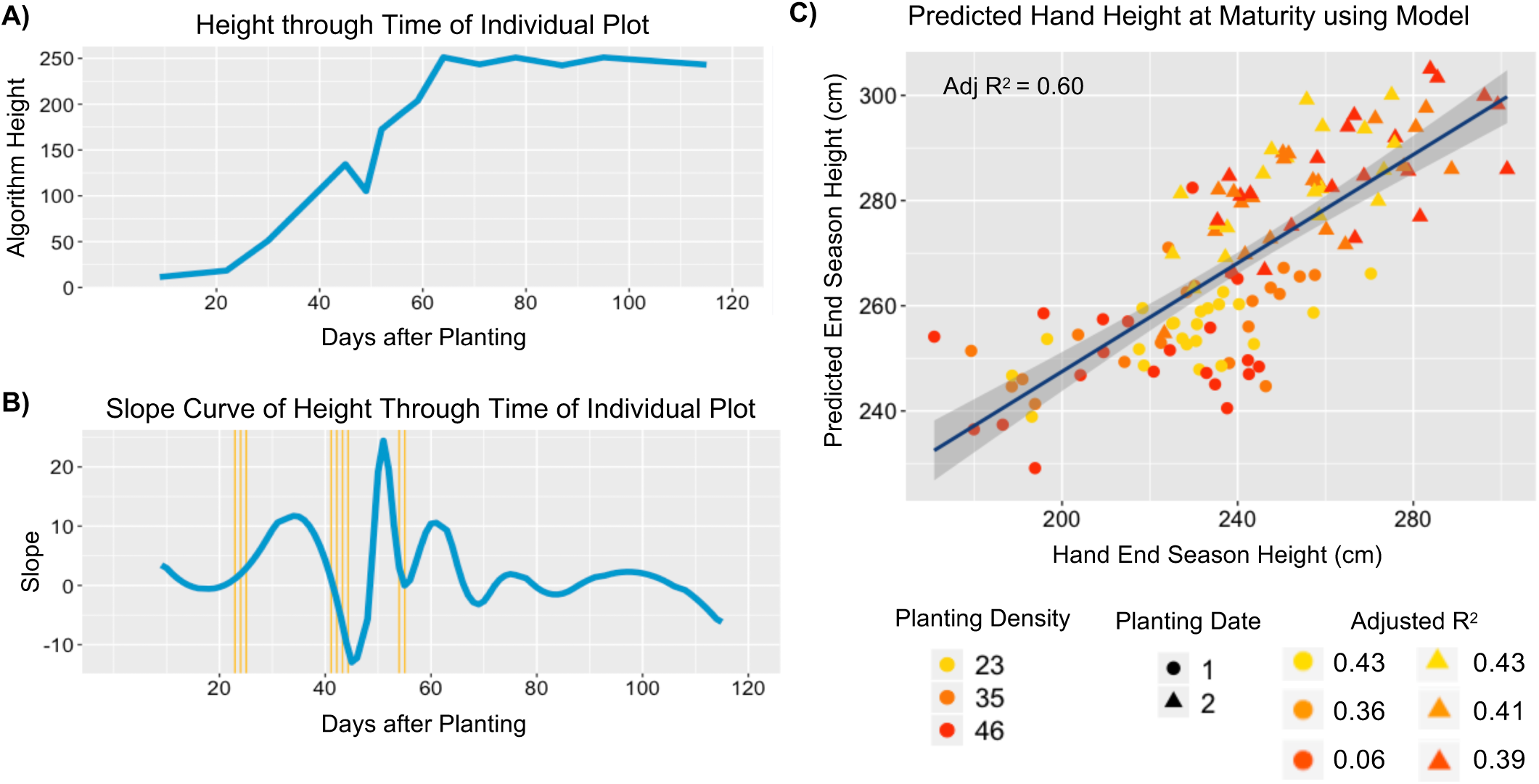
Predicting terminal height using rate of change in height at certain intervals. **A)** Height through time for an individual plot as recorded by the UAV. **B)** First derivative of the height through time spline curve in blue and the selected timepoints were the slope contributed most to the prediction of end-season height in yellow. **C)** Correlation of predicted end-season height on the test dataset utilizing a linear regression model derived from the training dataset based on slope values of selected timepoints and the hand-measured height values at maturity.

The slope curve was broken up into 100 equidistant timepoints for each plot and the contribution of each timepoint to the mean end-season height measured by hand was tested. A linear regression model was then created to predict end-season height based on the slope value for the timepoints that contributed more than 40% of the observed variation for terminal hand height.

The model was derived from ¼ of the data points and tested to predict the other ¾ of the plots. It contained nine timepoints, three of which were in the V2-V4 developmental stages, four in the V5-V7 stages and two in the V8-V10 stages. These timepoints all had biological significance as they fell upon areas of large inflection in the plant growth curve where a weather event or developmental time point caused a dramatic change in height. The first three timepoints which fall upon the V2-V4 developmental stage exhibited a large increase in height characteristic of early plant growth. The second set of three timepoints follow the storm that occurred around the V5-V7 developmental stage which caused a large degree of lodging just prior to their recovery. The last set of timepoints in the V8-V10 stages show a decline in rate of growth followed by a sudden increase. This decrease could be a consequence of weather variables such as temperature and moisture availability whereas the increase could be due to the emergence of tassels. These timepoints were crucial points in plant development where plant growth responded to certain conditions that in turn affected the end-season height obtained.

This model had a high ability to predict end-season mean plot PH_UAV_ (adjusted r-squared value of 0.63) and mean plot PH_R_ (adjusted r-squared value of 0.60; Figure 7C). When looking at the different treatments separately, we can see that the ability to predict height across planting dates increases for the late planting treatment and the magnitude of this difference increases with higher planting densities. This is likely due to the challenges in estimating mean plot height for plots that suffered lodging and the higher density treatments in the early planting date experienced the most lodging early in the season. These yield prediction results, although specific to this experiment, show the significant correlation that change in height from early season timepoints can have with terminal height and highlight the potential for developing tools to predict end-season height utilizing temporal PH measurements early in development. Maize plants uptake negligible amounts of nutrients such as nitrogen for the first weeks after emergence and then start to exponentially take up more nitrogen until the tasseling stage (Sharma and Bali, 2018). Being able to use height information gathered before the V10 developmental stage to predict end-season height and potentially stress can be a useful tool for guiding nutrient management practices early in development when plants are efficiently absorbing nutrients and can have a high response to fertilization.

## CONCLUSIONS

High-throughput collection of plant height measurements throughout the growing season can allow a better understanding of how different environments influence maize growth and can reveal important factors affecting crop productivity. We have developed a novel approach for extracting plot PH_UAV_ from individual plots that involves estimating the ground height individually for each plot and using that to calculate mean plot height. Overall, PH_R_ and PH_UAV_ measurements showed a high correlation (adj. r-square value of 0.96); however, correlations within plant height estimates collected on a single date are much lower as they are impacted by the low variability in height expressed on a single date. The reproducibility and quality of PH_R_ estimates were found to vary throughout development with lower correlations but smaller RMSEs were observed in early vegetative stages, and improved correlations but larger RMSEs were observed in later developmental stages. When assessing the error in generalizing mean PH_R_ for a plot by measuring only a subset of plants in a row, PH_R_ measurements were found to be greatly variable and impacted by the number and selection of plants sampled. Plot mean PH_R_ measurements obtained from different sets of plants within a row show a correlation and error across most timepoints comparable to those obtained when correlating plot mean PH_UAV_ measurements to plot mean PH_R_ measurements.

Implementing this pipeline to visualize height across different genotypes, planting densities and planting date treatments showed a strong ability to monitor biological variation for height across development in response to the environment. Differences in rate of growth between genotypes and planting dates as well as in lodging responses between planting densities were documented. Moreover, we saw that growth rates generated by PH measurements collected at multiple timepoints early in development can be useful in improving predictions of PH at the end of the season. Together, these findings show the utility of using UAVs for collecting PH data at high temporal resolution to track differences in growth and to predict an end-season trait early in development.

## Conflict of interest

The authors do not have any conflict of interest to declare.

## SUPPLEMENTAL MATERIAL

**Figure S1.** Field experimental layout for 2018 biological material.

**Figure S2.** Correlation of predicted end-season height on the test dataset.

**Table S1.** Adjusted r-square values and root mean square error for the linear correlation of various PH measurements.

**Table S2.** Analysis of variance of UAV-derived terminal plant height for 12 hybrids planted in a randomized block design with density and planting date treatments.

## SUPPLEMENTAL FIGURE LEGENDS

**Figure S1. Field experimental layout for 2018 biological material.**

**Figure S2. Correlation of predicted end-season height on the test dataset.** Predictions were done utilizing a linear regression model derived from the training dataset based on slope values of selected timepoints and the UAV derived height values at maturity.

**Table S1. Adjusted r-square values and root mean square error for the linear correlation of various PH measurements.** From left to right: replicated hand measurements obtained for 2 plants across 12 plots, means derived from the replicated hand measurements obtained for 2 plants of the same 12 plots, algorithm-derived plot mean height values to the respective hand-measured plot mean height of the same 12 plots, and model-derived plot mean height values compared to the respective hand-measured plot mean height of the same 12 plots.

**Table S2. Analysis of variance of UAV-derived terminal plant height for 12 hybrids planted in a randomized block design with density and planting date treatments.**

