## Supplemental Material for "UAV Based Imaging Platform for Monitoring Maize Growth Throughout Development"

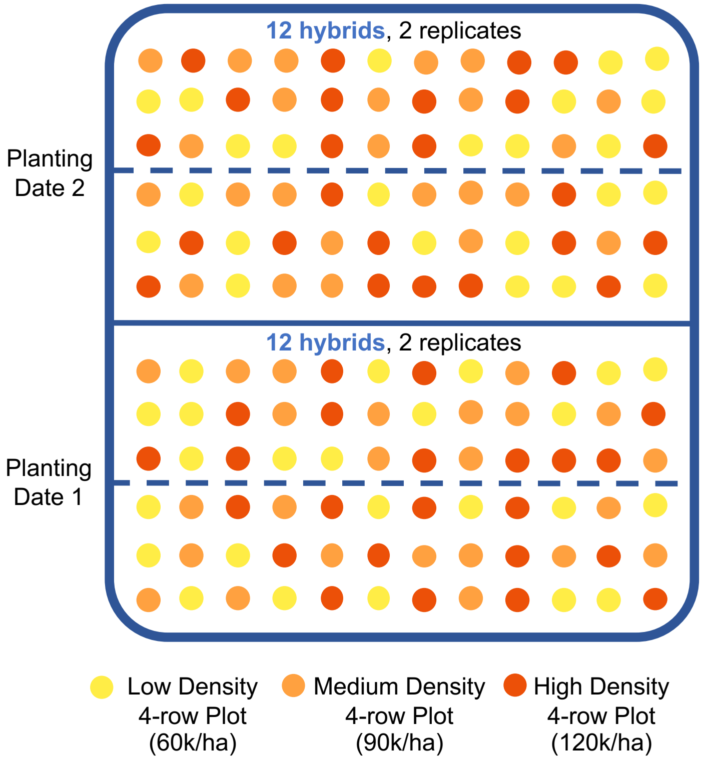


**Figure S1. Field experimental layout for 2018 biological material.**

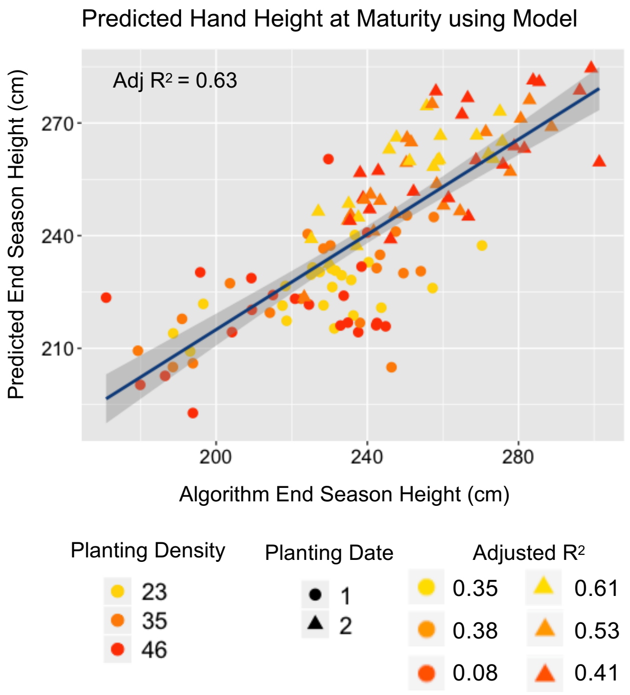


**Figure S2. Correlation of predicted end-season height on the test dataset.** Predictions were done utilizing a linear regression model derived from the training dataset based on slope values of selected timepoints and the UAV derived height values at maturity.

**Table S1. Adjusted r-square values and root mean square error for the linear correlation of various PH measurements.** From left to right: replicated hand measurements obtained for 2 plants across 12 plots, means derived from the replicated hand measurements obtained for 2 plants of the same 12 plots, algorithm-derived plot mean height values to the respective hand-measured plot mean height of the same 12 plots, and model-derived plot mean height values compared to the respective hand-measured plot mean height of the same 12 plots.

| | PH-R Replicates for Individual | | Plot Mean PH-UAV to PH-R for Plot Subset | | Plot Mean PH-R Replicates for Plot Subset | | Mean Plot PH-Model to PH-R for Plot Subset | |
|---|---|---|---|---|---|---|---|---|
| | **Adj R2** | **RMSE** | **Adj R2** | **RMSE** | **Adj R2** | **RMSE** | **Adj R2** | **RMSE** |
| 6/13/18 | 0.45 | 4.50 | 0.05 | 12.29 | 0.64 | 2.52 | 0.36 | 6.77 |
| 7/17/18 | 0.97 | 7.75 | 0.47 | 27.36 | 0.88 | 4.00 | 0.55 | 13.67 |
| 8/9/18 | 0.90 | 7.18 | 0.72 | 27.19 | 0.98 | 3.00 | 0.85 | 9.10 |

**Table S2. Analysis of variance of UAV-derived terminal plant height for 12 hybrids planted in a randomized block design with density and planting date treatments.**

| | Degrees of Freedom | Sum of Squares | Mean Sum of Squares | Percent of total variation |
|---|---|---|---|---|
| Planting Date | 1 | 46485 | 46485 | 43.37*** |
| Genotype | 11 | 11316 | 1029 | 10.56** |
| Density | 2 | 148 | 74 | 0.14 |
| Planting Date:Genotype | 11 | 4582 | 417 | 4.27 |
| Density:Genotype | 22 | 6218 | 283 | 5.80 |
| Residuals | 96 | 38443 | 418 | 35.86 |

* significant at $p$ =0.05; ** significant at $p$ =0.01; *** significant at $p$ =0.001; N.S., not significant.
